# A Rev-CBP80-eIF4AI complex drives Gag synthesis from the HIV-1 unspliced mRNA

**DOI:** 10.1101/313312

**Authors:** Daniela Toro-Ascuy, Bárbara Rojas-Araya, Francisco García-de-Gracia, Cecilia Rojas-Fuentes, Camila Pereira-Montecinos, Aracelly Gaete-Argel, Fernando Valiente-Echeverría, Théophile Ohlmann, Ricardo Soto-Rifo

## Abstract

Gag synthesis from the full-length unspliced mRNA is critical for the production of the viral progeny during human immunodeficiency virus type-1 (HIV-1) replication. While most spliced mRNAs follow the canonical gene expression pathway in which the recruitment of the nuclear cap-binding complex (CBC) and the exon junction complex (EJC) largely stimulates the rates of nuclear export and translation, the unspliced mRNA relies on the viral protein Rev to reach the cytoplasm and recruit the host translational machinery. Here, we confirm that Rev ensures high levels of Gag synthesis by driving nuclear export and translation of the unspliced mRNA. These functions of Rev are supported by the CBC subunit CBP80, which binds Rev and the unspliced mRNA in the nucleus and the cytoplasm. We also demonstrate that Rev interacts with the DEAD-box RNA helicase eIF4AI, which translocates to the nucleus and cooperates with Rev to promote Gag synthesis. Interestingly, molecular docking analyses revealed the assembly of a Rev-CBP80-eIF4AI complex that is organized around the Rev response element (RRE). Together, our results provide further evidence towards the understanding of the molecular mechanisms by which Rev drives Gag synthesis from the unspliced mRNA during HIV-1 replication.

## Introduction

Human Immunodeficiency Virus type-1 (HIV-1) gene expression is a complex process that leads to the synthesis of fifteen proteins from one single primary transcript (1,2). Once the proviral DNA has been integrated into the host cell genome, the RNA polymerase II drives the synthesis of a 9-kb, capped and polyadenylated pre-mRNA that undergoes alternative splicing generating more than 100 different transcripts classified into three main populations (3,4). The so-called 2-kb multiply spliced transcripts code for the key regulatory proteins Tat and Rev and the accessory protein Nef and are the dominant viral mRNA species at early stages of viral gene expression (1,2,5). Unlike cellular mRNAs or the 2-kb transcripts, which are spliced to completion before they exit the nucleus, HIV-1 and other complex retroviruses produce an important fraction of viral transcripts that remain incompletely spliced (2,6). These 4-kb transcripts are expressed during the intermediate phase of gene expression and are used for the synthesis of the envelope glycoprotein (Env) and the accessory proteins Vif, Vpr and Vpu (2,6). Finally, the full-length 9-kb pre-mRNA in its unspliced form also reaches the cytoplasm to be used as an mRNA template during the late stages of viral gene expression for the synthesis of the major structural proteins Gag and Gag-Pol (1,2,6).

Gene expression in eukaryotic cells occurs through the intricate connection of different processes including transcription, splicing, nuclear export, translation and mRNA decay and is regulated by the specific recruitment of nuclear proteins that together form the messenger ribonucleoprotein (mRNP) complex (7-9). As such, the early binding of the nuclear cap-binding complex (CBC) to the 5´-end cap structure and the splicing-dependent recruitment of nuclear proteins such as the exon-junction complex (EJC) onto the mRNA has been shown to increase the rates of nuclear export and translation of spliced transcripts (10-20). Eukaryotic cells have also evolved quality control mechanisms ensuring that only properly processed mRNAs reach the cytoplasm and are decoded by the translational machinery. These mechanisms include the EJC-dependent degradation of transcripts containing premature stop codons through nonsense-mediated decay (NMD) or the NXF1-dependent nuclear retention of unspliced transcripts mediated by the nucleoporin Tpr (21- 25). Consistent with these cellular quality control mechanisms, it has been widely reported that viral intron-containing transcripts including the 4-kb and the 9-kb mRNA produced during HIV-1 replication are retained and degraded in the host cell nucleus unless the viral protein Rev is present (26-30). As such, Rev has been proposed to promote HIV-1 gene expression from its target transcripts by i) avoiding mRNA degradation (26,31); ii) promoting nuclear export (31-33) or by iii) promoting translation (34,35). In this study, we developed Rev mutant proviruses and confirmed that Rev is required for both nuclear export and translation of the HIV-1 unspliced mRNA. Interestingly, we show that the nuclear cap-binding complex subunit CBP80 and the translation initiation factor eIF4AI associate with Rev and the unspliced mRNA to promote Gag synthesis. Molecular docking analyses suggest the assembly of a Rev-CBP80-eIF4AI complex that is reorganized when Rev binds to the RRE. Together, our work provides further insights into the molecular mechanism by which Rev drives Gag synthesis from the HIV-1 unspliced mRNA.

## Materials and methods

### DNA constructs

The pNL4.3 and pNL4.3R proviruses were previously described (36,37). These vectors were digested with NheI and subjected to a 20 min polymerization reaction at 72 ΰC using the Phusion® High-Fidelity DNA polymerase (New England Biolabs) in order to create a frameshift that generates a premature stop codon within the *env* gene. The resulting vectors were ligated with the T4 DNA ligase and transformed into *E.coli* DH5α. To create the pNL4.3-ΔRev and pNL4.3R-ΔRev vectors, the above vectors were digested with BamHI and subjected to the same polymerization/ligation reaction to create a frameshift within the *rev* gene previously shown to abolish expression of a functional protein (28). The pCMV-NL4.3R and pCMV-NL4.3R-ΔRev vectors were obtained by replacing the FspAI/BssHII fragment of the corresponding vector by the CMV IE promoter amplified from the pCIneo vector (Promega) as we previously reported (38). The pCDNA-Flag-Rev vector was previously described (22). pCDNA-d2EGFP vector was generated by inserting the d2EGFP ORF into pCDNA3.1 (Life Technologies). pCDNA HIV-1 5´-UTR and pCDNA β-globin 5´-UTR were previously described (39). The dl HIV-1 IRES vector was previously described (40). The pCIneo-HA-eIF4GI,-eIF4A and -eIF4E were previously described (41). The pCMV-myc-eIF4E and CBP80 were previously described (42).

### Cell culture and DNA transfection

HeLa cells and human microglia (C20 cells)(43) were maintained in DMEM (Life Technologies) supplemented with 10% FBS (Hyclone) and antibiotics (Hyclone) at 37 °C and a 5% CO_2_ atmosphere. H9 T-lymphocytes (44) and THP-1_ATCC_ monocytes (45) were maintained in RPMI 1640 (Life Technologies) supplemented with 10% FBS (Hyclone) and antibiotics (Hyclone) at 37 °C and a 5% CO_2_ atmosphere. Cells were transfected using linear PEI ~25.000 Da (Polysciences) prepared as described previously (46). Cells were transfected using a ratio μg DNA/μl PEI of 1/15.

### Analysis of Renilla and firefly activities

Renilla activity was determined using the Renilla Reporter Assay System (Promega) and Renilla/firefly activities were determined using the Dual Luciferase Reporter Assay System (Promega) in a GloMax® 96 microplate luminometer (Promega).

### Western blot

Cells extracts from transfected cells were prepared by lysis with RIPA buffer and 20 μg of total protein were subjected to 10% SDS-PAGE and transferred to an Amersham Hybond™-P membrane (GE Healthcare). Membranes were incubated with an HIV-1 p24 monoclonal antibody diluted to 1/1000 (47), a rabbit anti-Flag antibody (Sigma-Aldrich) diluted to 1/1000 or and HRP-conjugated anti-actin antibody (Santa Cruz Biotechnologies) diluted to 1/750. Upon incubation with the corresponding HRP-conjugated secondary antibody (Santa Cruz Biotechnologies) diluted to 1/1000, membranes were revealed with the ECL substrate (Cyanagene) using a C-Digit digital scanner (Li-Cor).

### RNA extraction and RT-qPCR

Cytoplasmic RNA extraction and RT-qPCR from cytoplasmic RNA were performed essentially as recently described (48). Briefly, cells were washed intensively with PBS, recovered with PBS-EDTA 10 mM and lysed for 1-2 min at room temperature with 200 μl of buffer [10 mM Tris-HCl pH=8.0, 10 mM NaCl, 3 mM MgCl2, 1 mM DTT, 0.5% NP40 and 2 mM of vanadyl-ribonucleoside complex (VRC) (New England Biolabs)]. Cell lysates were centrifuged at 5000 rpm for 5 min at 4 °C and supernatant containing the cytoplasmic fraction were recovered and RNA extraction was carried out by adding 1 ml of TRIzol^™^ (Thermo Fisher) as indicated by the manufacturer. Cytoplasmic RNAs (1 μg) were reverse-transcribed using the High Capacity RNA-to-cDNA Master Mix (Life Technologies). For quantitative PCR, a 20 μl reaction mix was prepared with 5 μl of template cDNAs (previously diluted to 1/10), 10 μl of FastStart Universal SYBR Green Master (Rox) (Roche), 0.2 μM of sense and antisense primers and subjected to amplification using the Rotorgen fluorescence thermocycler (Qiagen). The GAPDH housekeeping gene was amplified in parallel to serve as a control reference. Relative copy numbers of Renilla luciferase cDNAs were compared to GAPDH or 18S rRNA using (Inline) (where x correspond to the experimentally calculated amplification efficiency of each primer couple).

### Fluorescent in situ hybridization, immunofluorescence and confocal microscopy

RNA FISH was carried out as we recently described (48). Briefly, HeLa cells were cultured in a 12 well plate with covers slide (Nunc^™^) and maintained and transfected with 1 μg of pNL4.3 or 1 μg of the corresponding HA vectors as indicated above. At 24 hpt, cells were washed twice with 1X PBS and fixed for 10 min at room temperature with 4% paraformaldehyde. Cells were subsequently permeabilized for 5 min at room temperature with 0.2% Triton X-100 and hybridized overnight at 37° C in 200 μl of hybridization mix (10% dextran sulfate, 2 mM VRC, 0.02% RNase-free BSA, 50% formamide, 300 μg tRNA and 120 ng of 11-digoxigenin-UTP probes) in a humid chamber. Cells were washed with 0.2X SSC/50% formamide during 30 min at 50° C and then incubated three times with antibody dilution buffer (2X SSC, 8% formamide, 2 mM vanadyl-ribonucleoside complex, 0.02% RNase-free BSA). Mouse anti-digoxin and rabbit anti-HA (Sigma Aldrich) primary antibodies diluted to 1/100 in antibody dilution buffer were added for 2 h at room temperature. After three washes with antibody dilution buffer, cells were incubated for 90 min at room temperature with anti-mouse Alexa 488 and anti-rabbit Alexa 565 antibodies (Molecular Probes) diluted at 1/1000. Cells were washed three times in wash buffer (2X SSC, 8% formamide, 2 mM vanadyl-ribonucleoside complex), twice with 1X PBS, incubated with Hoescht (Life Technologies) diluted to 1/10000, for 5 min at room temperature, washed three times with 1X PBS, three times with water and mounted with Fluoromount (Life Technologies). Images were obtained with a TCS SP8 Confocal Microscope (Leica Microsystems) and images were processed using FIJI/ImageJ (NIH).

### Proximity ligation assay (PLA)

PLA (49), was carried out using the DUOLINK II In Situ kit (Sigma-Aldrich) and PLA probe anti-mouse minus and PLA probe anti-rabbit plus (Sigma-Aldrich) as we have previously described (38,50). Briefly, PFA-fixed HeLa cells were pre-incubated with blocking agent for 30 min at room temperature. Primary antibodies were added at a dilution of 1:100 (mouse anti-HA, Santa Cruz Biotechnologies) and 1:200 (rabbit anti-Flag, Sigma-Aldrich) in 40 μl DUOLINK antibody diluent and incubated at 37°C for 1 h. Samples were washed three times with PBS for 5 min each and secondary antibodies (DUOLINK anti-rabbit PLA-plus probe and DUOLINK anti-mouse PLA-minus probe) were added and incubated at 37°C for 1 h. Ligation and amplification reactions were performed following the same protocol described in (50). Samples were incubated with DAPI (0.3 μg/ml in PBS) (Life Technologies) for 1 min at room temperature, washed three times with PBS, three times with water and mounted with fluoromount (Sigma Aldrich). Images were obtained with a TCS SP8 Confocal Microscope (Leica Microsystems) and images were processed using FIJI/ImageJ (NIH).

### In situ hybridization coupled to PLA (ISH-PLA)

The ISH-PLA protocol was developed by mixing the RNA-FISH and PLA protocols described above. Briefly, PFA-fixed Hela cells growing on coverslips were permeabilized for 5 min at RT with 0.2% Triton X-100 and hybridized overnight at 37°C in 200 μl of hybridization mix (10% dextran sulphate, 2mM vanadyl–ribonucleoside complex, 0.02% RNase-free bovine serum albumin, 50% formamide, 300 mg of tRNA and 120 ng of 11-digoxigenin-UTP probes) in a humid chamber. Cells were washed with 0.2xSSC/50% formamide during 30 min at 50°C and then incubated with blocking agent for 30 min to room temperature. Then incubated three times with antibody dilution buffer (2xSSC, 8%formamide, 2mM vanadyl–ribonucleoside complex and 0.02% RNase-free bovine serum albumin). Mouse anti-digoxin and rabbit anti-protein of interest primary antibodies diluted to 1/100 in antibody dilution buffer were added for 2 h at room temperature. After three washes with antibody dilution buffer and two washes with PBS for 5 min each, then secondary antibodies (DUOLINK anti-rabbit PLA-plus probe, DUOLINK anti-mouse PLA-minus probe) were added and incubated at 37°C for 1 h. Then, the ligation and amplification reaction were performed following the same protocol described above. Thereafter, the covers were incubated with a solution of DAPI (Life Technologies) (0.3 μg/ml in PBS) for 1 min at room temperature, washed three times with PBS, three times with water and mounted with fluoromount aqueous mounting medium (Sigma Aldrich). Images were obtained with a TCS SP8 Confocal Microscope (Leica Microsystems) and the Images were processed using FIJI/ImageJ (NIH) (Supplementary Fig. 2A).

### Bioinformatic analyses

Molecular docking was performed using previously described structures of CBP80 (PDB: 3FEY) (51), eIF4AI (PDB: 2ZU6) (52), Rev monomer (PDB: 2X7L) (53), Rev dimer (PDB: 3LPH) (54) and Rev dimer bound to the RRE (PDB: 4PMI)(55). Since the structure of the Rev monomer contains the L12S/L60R mutations (54), we proceed with a structural analysis between the dimer and the monomer using the SALIGN module of MODELLER, version 9.13 (56). Structure visualization was performed with Visual Molecular Dynamics (VMD) (57). The local quality of the wild type Rev dimer was estimated with ANOLEA (Anolea, atomic mean force potential) (58) for energy evaluation and with PROCHECK for stereochemistry evaluation (59). Subsequently, an energy minimization of 5000 steps was performed using the Steepest Descent (SD) method. For protein-protein interactions, the crystallized structures were treated by the SD method described above prior processing and refinement of the model. Macromolecular complexes were obtained with PATCHDOCK (60), which analyses separately the surface of both proteins and generates geometric patterns depending on the shape complementary of soft molecular surfaces in order to generate the best starting candidate solution. Refinement of the side-chain flexibility, rigid body optimization, scoring of docking candidates and ranking were performed with FIREDOCK (61). Macromolecular complexes were selected based on their energetic characteristics and the non-covalent interactions were determined with the Discovery Studio Visualizer software.

## Results

### Rev promotes nuclear export and translation of the unspliced mRNA

The viral protein Rev promotes gene expression from the unspliced transcript by acting at the post-transcriptional level but the precise mechanism by which this occurs remains unclear (62-64). This prompted us to conduct a study aimed to gain further insights into the function of Rev on HIV-1 gene expression during viral replication. Thus, we used the pNL4.3 proviral DNA to introduce a frameshift within the *rev* gene previously shown to abolish the expression of a functional protein (28). Consistent with a critical role of Rev in the post-transcriptional regulation of the unspliced mRNA, we observed that Gag synthesis was abolished in the absence of Rev and restored upon Rev expression *in trans* (Fig. 1A). In agreement with several previous reports (31-33), we observed that most of the unspliced mRNA is retained in the nucleus in the absence of Rev (Fig. 1B, compare wild type and ΔRev). The cytoplasmic signal of the unspliced mRNA was recovered when Rev was expressed *in trans* (Fig. 1B, see ΔRev + Flag-Rev), which is consistent with an important role of Rev in nuclear export. However, it has been reported that Rev is also important for translation of the unspliced mRNA during viral replication (34,35). Thus, in order to quantify the impact of Rev on gene expression from the unspliced mRNA we used the pNL4.3R reporter provirus to generate a ΔRev version as described above (see materials and methods). Transfection of pNL4.3R-ΔRev in HeLa cells, T-cells (H9 cells), monocytes (THP-1 cells) or human microglia (C20 cells) resulted in very low levels of Gag-Renilla expression when compared to the wild type provirus indicating that our reporter proviruses can be used to quantify the effects of Rev on gene expression from the unspliced mRNA (Supplementary Fig. 1A). Transfection of the pNL4.3R-ΔRev together with a Flag-Rev expressing vector restored Gag synthesis to the wild type levels indicating that the defects in Gag-Renilla expression observed were exclusively due to the absence of Rev (Supplementary Fig. 1B). Thus, we used the pNL4.3R-wt and -ΔRev proviruses to quantify the effects of Rev on Gag synthesis, cytoplasmic levels of the unspliced mRNA and its translational efficiency in HeLa cells as we have previously reported (37,38,41,48,65,66). As observed above, Gag production was almost abolished in the absence of Rev (Fig. 1C, left panel). Consistent with its previously described role in nuclear export of the unspliced transcript (26, 31-33), we observed that the cytoplasmic levels of the unspliced mRNA were reduced by 3-fold in the absence of Rev (Fig. 1C, middle panel). Although these results differ from those presented in Fig. 1B in which no genomic RNA could be detected in the cytoplasm, it has been proposed that an important fraction of the unspliced transcript reach the cytoplasm in the absence of Rev but remains trapped into a ribonucleoprotein complex inaccessible to the probes used during *in situ* hybridization (31). It should be mentioned that our cytoplasmic fractions were devoid of pre-GADPH mRNA discarding any contamination with nuclear RNA (Supplementary Fig. 1C). However, this important decrease in the cytoplasmic levels of the unspliced transcript does not account for the dramatic reduction in Gag synthesis (>100-fold) indicating that most of the unspliced mRNAs that reach the cytoplasm in the absence of Rev are not translated (Fig. 1C, right panel). Together, these data confirm that the function of Rev during viral replication is not restricted to nuclear export since it is also critical for translation.

**Figure 1.**
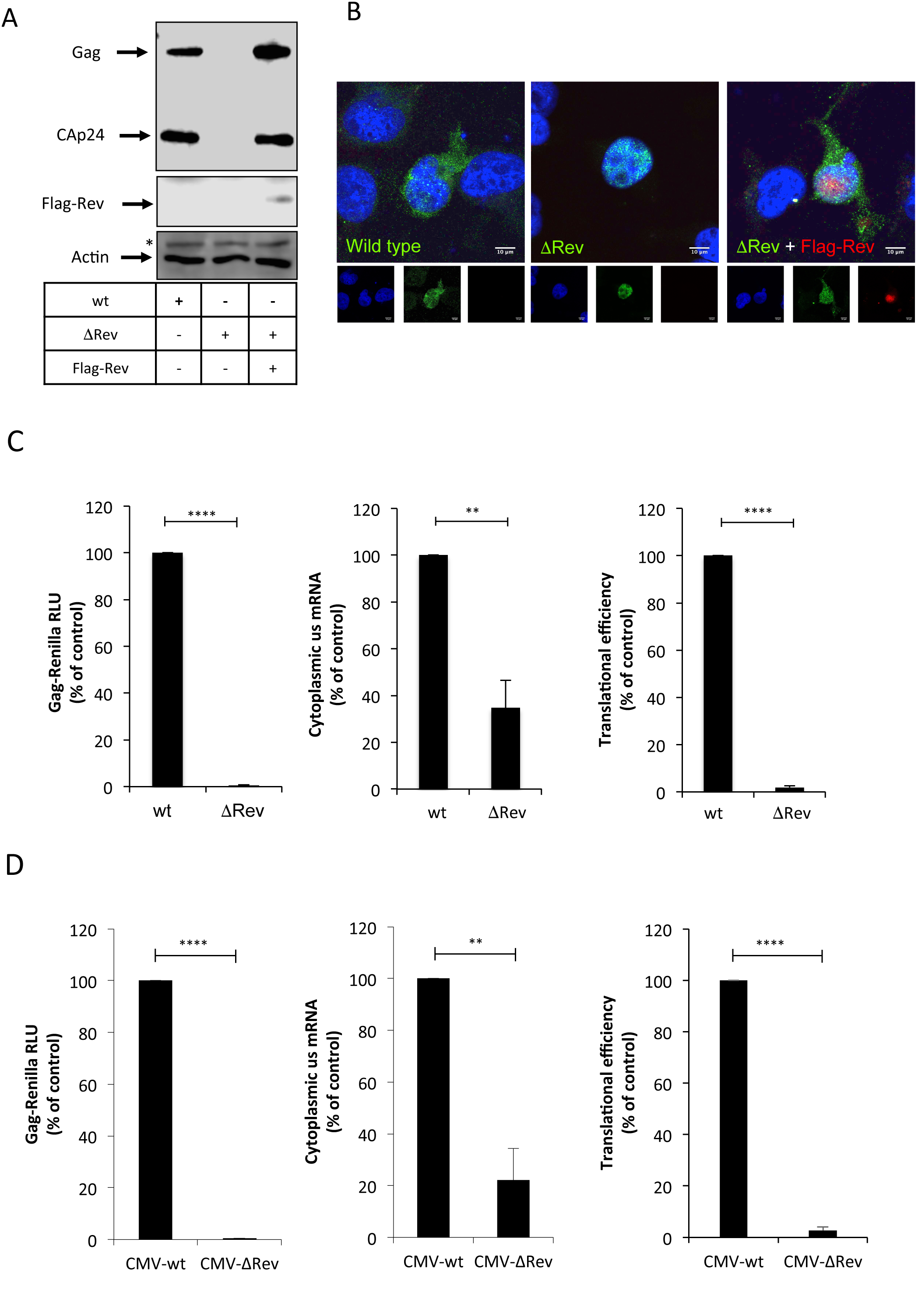
HIV-1 Rev promotes nuclear export and translation of the unspliced mRNA. A) HeLa cells were transfected with 0.3 μg of pNL4.3-wt (wt), pNL4.3-ΔRev (ΔRev) or pNL4.3-ΔRev together with 0.1 μg of the pCDNA-Flag-Rev vector as described in materials and methods (pCDNA-d2EGFP was used as a control when Flag-Rev was not included). At 24 hpt, cell extracts were used to detect Gag and Flag-Rev by Western blot. Actin was used as a loading control. ^*^ Denotes an unspecific band detected with the anti-actin-HRP antibody. B) HeLa cells were transfected as above and were subjected to RNA FISH and laser scan confocal microscopy analyzes as described in materials and methods. The unspliced mRNA is showed in green and Flag-Rev in red. Scale bar 10 μm. C) HeLa cells were transfected with 0.3 μg of pNL4.3R (wt) or pNL4.3R-ΔRev (ΔRev) proviruses as described in materials and methods. At 24 hpt, cell extracts were prepared for Gag-Renilla activity measurement and for cytoplasmic RNA extraction and RT-qPCR analyzes. Results for Gag synthesis (left panel), cytoplasmic unspliced mRNA (middle panel) and translational efficiency (right panel) were normalized to the wild type provirus (arbitrary set to 100%) and presented as the mean +/- SD of three independent experiments (^**^*p*<0.01; ^****^*p*<0.0001, t-test). D) HeLa cells were transfected with 0.3 μg of pCMV-NL4.3R-wt (CMV-wt) or pCMV-NL4.3R-ΔRev (CMV-ΔRev) proviruses as described in materials and methods. At 24 hpt, cell extracts were prepared for Gag-Renilla activity measurement and for cytoplasmic RNA extraction and RT-qPCR analyzes. Results for Gag synthesis (left panel), cytoplasmic unspliced mRNA (middle panel) and translational efficiency (right panel) were normalized to the wild type provirus (arbitrary set to 100%) and presented as the mean +/- SD of three independent experiments (^**^*p*<0.01; ^****^*p*<0.0001, t-test).

While the role of Rev in nuclear export has been largely characterized (62), the molecular mechanism by which Rev promotes translation of the unspliced transcript is not very well understood (67). Interestingly, it was shown that Rev was able to promote translation of a reporter RNA by an unknown mechanism involving the binding to an RNA motif (A-loop) present within SL1 of the unspliced mRNA 5´-UTR (63,68). Consistent with this previous work, we observed that expression of a Renilla luciferase-based monocistronic vector harboring the 5´-UTR of the unspliced transcript (but not that of the human β-globin) was stimulated up to 2-fold in the presence of Rev (Supplementary Fig. 1D). Such a stimulation was not observed when IRES-driven translation was analyzed with a previously described bicistronic vector (40), suggesting that the effect of Rev on the 5´-UTR is exerted at the level of cap-dependent translation (Supplementary Fig. 1E).

Although the 2-fold stimulation observed with the reporter construct is consistent with the previous report (63), it does not account for the strong dependence of Rev for unspliced mRNA translation observed in the context of a full-length provirus (Fig. 1A and 1C), suggesting that the 5´-UTR is not the only molecular determinant involved in Rev-mediated translation of the unspliced mRNA. In order to confirm this hypothesis, we constructed a ΔRev version of the CMV-pNL4.3R vector, a reporter provirus lacking the 5´-LTR and most of the 5´-UTR but containing the CMV IE promoter (38). This proviral DNA is expected to produce an unspliced mRNA that only contains the last 79 nucleotides of the wild type 5´-UTR, therefore lacking the Rev binding site previously described in the A-loop of SL1. We transfected the CMV-pNL4.3R and CMV-pNL4.3R-ΔRev vectors and analyzed the role of Rev in Gag synthesis, cytoplasmic levels of the unspliced mRNA and translation as described above. Interestingly, we observed a strong dependence for Rev in translational efficiency of the unspliced transcript regardless of whether the entire 5´-UTR was driving ribosome recruitment or not (Fig. 1D). Together, these data suggest that the previously described Rev-binding site present within the 5´-UTR is not the major molecular determinant involved in the translational stimulation mediated by Rev in the context of viral replication. Given the fact that our CMV-pNL4.3R provirus also lacks major determinants required for IRES-driven translation (40), these data also suggest that Rev promotes cap-dependent translation.

### The CBC subunit CBP80 interacts with Rev and promotes nuclear export and translation of the unspliced mRNA

From data presented above, it seems that Rev promotes cap-dependent translation of the unspliced mRNA. Interestingly, it was proposed that cap-dependent CBC-driven translation could ensure Gag synthesis during an HIV-induced inhibition of eIF4E activity (69). In agreement with this idea, we have previously shown that the cytoplasmic cap-binding protein eIF4E is excluded from a translation initiation mRNP containing the HIV-1 unspliced mRNA together with the RNA helicase DDX3 and translation initiation factors eIF4GI and PABPC1 (48). Thus, in order to determine whether the unspliced mRNA is associated to the CBC or eIF4E, we developed a protocol based on *in situ* hybridization of digoxin-labeled probes directed to the unspliced mRNA coupled to the proximity ligation assay (ISH-PLA) in order to determine and quantify unspliced mRNA-protein interactions (Supplementary Fig. 2A and materials and methods). By using our ISH-PLA protocol, we observed that the unspliced mRNA preferentially associates with the CBC subunit CBP80 rather than eIF4E (Fig. 2A). Interestingly, despite the fact that most of the CBP80 signal was observed in the nucleus in RNA FISH-IF experiments performed in parallel (Supplementary Fig. 2B), we observed that the interaction between the unspliced mRNA and the CBC subunit occurs predominantly in the cytoplasm suggesting that the unspliced mRNA remains associated to the CBC upon nuclear export. Signal intensity quantifications from the RNA FISH experiments performed in parallel revealed no differences in myc-tagged protein expression (Supplementary Fig. 2C). To further investigate the role of CBP80 and eIF4E on gene expression from the unspliced mRNA, we independently overexpressed both proteins in HeLa cells and analyzed Gag synthesis, cytoplasmic unspliced mRNA levels and translational efficiency as described above. Consistent with a preferential association of the unspliced mRNA with CBP80, we observed that overexpression of the CBC subunit but not eIF4E results in a marked increase in Gag synthesis, which was due to an increase in both cytoplasmic accumulation and translation of the unspliced mRNA (Fig. 2B). Interestingly, CBP80 overexpression only resulted in marginal (2-fold) stimulation of the unspliced mRNA in the cytoplasm when the ΔRev provirus was used suggesting that CBP80 cooperates with Rev during the post-transcriptional control of the unspliced mRNA (Fig. 2C). Of note, Gag-Renilla activity from the wild type provirus was largely higher than that observed from the ΔRev provirus consistent with data presented in Fig. 1 (data not shown). Consistent with the dependence on Rev for CBP80 function, we observed that CBP80 overexpression has marginal effects on protein synthesis from the Rev-independent nef mRNA (Supplementary Fig. 2D).

Since it was previously shown that CBP80 interacts with Rev *in vitro* (70), we wanted to evaluate whether this interaction also occurs in cells. Thus, we performed PLA and observed that Flag-tagged Rev interacts with both endogenous and V5-tagged CBP80 (Supplementary Fig. 2E and Fig. 2D).

Together, these results suggest that the unspliced mRNA is preferentially associated to the CBC and the CBC subunit CBP80 interacts and cooperates with Rev to promote nuclear export and translation of this viral transcript.

### DEAD-box helicase eIF4AI interacts with Rev and promotes Gag synthesis from the unspliced mRNA

Having determined that the unspliced mRNA is preferentially associated with CBP80 and that this CBC subunit interacts with Rev, we were interested in identify additional translation initiation factors interacting with Rev that could be involved in Gag synthesis. Although the CBP20/80-dependent translation initiation factor (CTIF) was shown to be important for CBC-dependent translation (71), we observed that CTIF is rather a potent inhibitor of Gag synthesis (García-de-Gracia *et al*, manuscript in preparation). Thus, we reasoned that Rev and CBP80 form an mRNP different from the canonical CBC, which is important for unspliced mRNA nuclear export and translation. Despite we showed that eIF4E seems not relevant for unspliced mRNA translation (Fig. 2A), we looked whether additional components of eIF4F such as eIF4GI and eIF4AI were associated with Rev.

**Figure 2:**
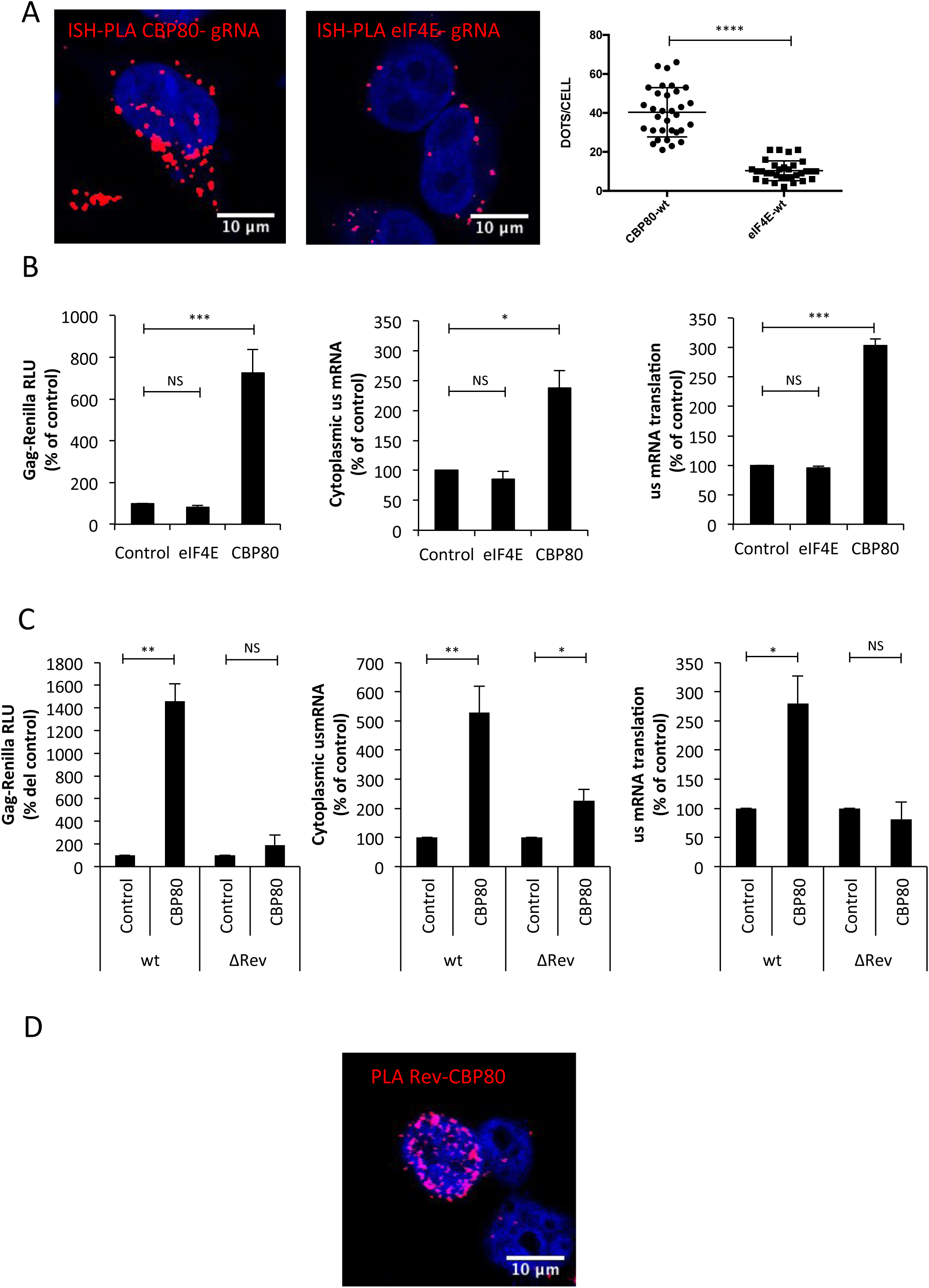
CBP80 cooperates with the functions of Rev on Gag synthesis from the unspliced mRNA. A) HeLa cells were transfected with 1 μg pNL4.3-wt together with 1 μg pCMV-myc-CBP80 or 1 μg pCMV-myc-eIF4E. At 24 hpt, the interaction between the unspliced mRNA and the myc-tagged protein was analyzed by the ISH-PLA protocol described in material and methods. Red dots indicate the interactions between the unspliced mRNA and the corresponding myc-tagged protein. Scale bar 10 μm. A quantification of the dots/cell in each condition is presented on the right (*p*<0.0001, Mann-Whitney test). B) HeLa cells were transfected with 0.3 μg of pNL4.3R or pNL4.3R ∆Rev proviruses together with 1 μg of pCMV-myc-CBP80 or pCMV-myc-eIF4E (pCVM-myc-d2EGFP was used as control). At 24 hpt, cell extracts were prepared for Gag-Renilla activity measurement and for cytoplasmic RNA extraction and RT-qPCR analyzes. Results for Gag synthesis (left panel), cytoplasmic unspliced mRNA (middle panel) and translational efficiency (right panel) were normalized to the wild type provirus (arbitrary set to 100%) and presented as the mean +/- SD of three independent experiments (^*^*p*<0.05; ^***^*p*<0.001 and NS; non-significant, t-test). C) HeLa cells were transfected with 0.3 μg of pNL4.3R or pNL4.3R ∆Rev proviruses together with 1 μg of pCMV-myc-CBP80 (pCVM-myc-d2EGFP was used as control) as described in materials and methods. At 24 hpt, cell extracts were prepared for Gag-Renilla activity measurement and for cytoplasmic RNA extraction and RT-qPCR analyzes. Results for Gag synthesis (left panel), cytoplasmic unspliced mRNA (middle panel) and translational efficiency (right panel) were normalized to the wild type provirus (arbitrary set to 100%) and presented as the mean +/- SD of three independent experiments. (^*^*p*<0.05; ^**^*p*<0.01 and *NS;* non-significant, t-test). D) HeLa cells were transfected with 1 μg pCDNA-Flag-Rev and 1 μg pCDNA-V5-CBP80. At 24 hpt, the interaction between Flag-Rev and V5-CBp80 was analyzed by PLA. Red dots indicate interactions between both proteins. Scale bar 10 μm.

Indeed, eIF4GI was shown to interact with CBP80 and thus, we supposed that it could be recruited to the Rev-CBP80 complex (72). Interestingly, our PLA using HA-tagged versions of eIF4E, 4GI and 4AI together with Flag-tagged Rev revealed that Rev forms nuclear and cytoplasmic complexes with eIF4AI and at a much lesser extent with eIF4E and eIF4GI (Figs. 3A and 3B). It should be mentioned that IF experiments performed in parallel revealed no differences in the intensity signals amongst HA-tagged eIFs and Flag-Rev indicating that the increased number of interactions observed between eIF4AI and Rev were not due to differences in the ectopic expression levels of the proteins (Supplementary Fig. 3A). From these data, it could be speculated that Rev recruits eIF4AI to the unspliced mRNA in order to promote translation. Thus, in order to evaluate the involvement of eIF4AI in Rev activity, we overexpressed the RNA helicase and analyzed its impact on Gag synthesis, cytoplasmic unspliced mRNA and translation using our wild type and ΔRev reporter proviruses (Fig. 3C). As expected, Gag-Renilla activity from the wild type provirus was much higher than that observed from the ΔRev provirus (data not shown). Interestingly, we observed that eIF4AI overexpression results in a 2- and 5-fold increase in Gag synthesis from the wild type and ΔRev reporter proviruses, respectively (Fig. 3C, left panel). Surprisingly, analysis of the cytoplasmic unspliced mRNA levels upon eIF4AI overexpression revealed a 2- to 3-fold increase for the wild type provirus with no effects on the ΔRev provirus (Fig. 3C, middle panel). More strikingly, we observed that translation from the ΔRev provirus was stimulated up to 6-fold by eIF4AI overexpression with no effects on translation from the wild type provirus (Fig. 3C, right panel). These results suggest that the presence of Rev will determine the process (nuclear export or translation) by which ectopically expressed eIF4AI will promote Gag synthesis from the unspliced mRNA.

**Figure 3:**
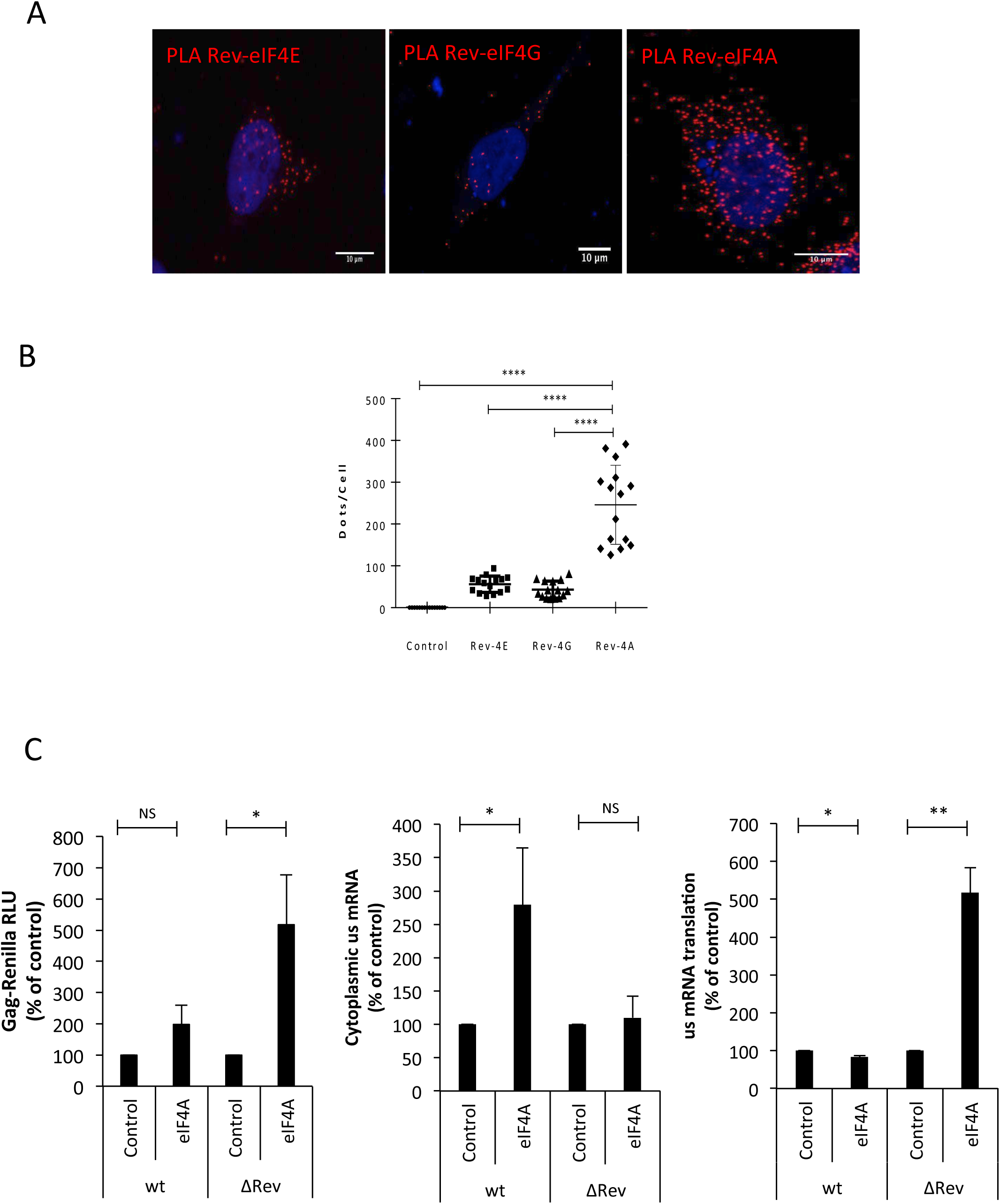
DEAD-box RNA helicase eIF4AI interacts with Rev and promotes Gag synthesis from the unspliced mRNA. A) HeLa cells were transfected with 1 μg pCDNA-Flag-Rev together with 1 μg pCIneo-HA-eIF4E, pCIneo-HA-eIF4G or pCIneo-HA-eIF4A. At 24 hpt, the interaction between Flag- and HA-tagged eIFs was analyzed by PLA. Red dots indicate interactions between Flag-Rev and the corresponding HA-tagged protein. Scale bar 10 μm B) Dots/cell for Rev-4E, Rev-4G and Rev-4A were quantified using ImageJ (*p*<0.0001, Mann-Whitney test). C) HeLa cells were transfected with 0.3 μg of pNL4.3R-wt or pNL4.3R-ΔRev together with 1 μg of the pCIneo-HA-eIF4A vector as described in materials and methods (pCIneo-HA-d2EGFP was used as a control). At 24 hpt, cell extracts were prepared for Gag-Renilla activity measurement and for cytoplasmic RNA extraction and RT-qPCR analyzes. Results for Gag synthesis (left panel), cytoplasmic unspliced mRNA (middle panel) and translational efficiency (right panel) were normalized to the wild type provirus (arbitrary set to 100%) and presented as the mean +/- SD of three independent experiments (^*^*p*<0.05; ^**^*p*<0.01 and *NS;* non-significant, t-test).

### Rev regulates the association of CBP80 and eIF4AI to the unspliced mRNA

Giving the fact that the presence of Rev modulates the activity of CBP80 and eIF4AI on Gag synthesis, we wanted to determine whether Rev was involved in the recruitment of these cellular proteins to the unspliced mRNA. For this, we quantified the interaction between the unspliced mRNA and CBP80 or eIF4AI in the presence and absence of Rev using our ISH-PLA protocol. Interestingly, we observed that the interactions between CBP80 and the unspliced mRNA were reduced in the ΔRev provirus but were restored upon expression of Rev *in trans* suggesting that Rev favors and/or stabilizes the association between CBP80 and the unspliced mRNA (Fig. 4A, left panel). Interestingly, we observed that Rev is necessary to maintain the CBP80-unspliced mRNA interaction in the cytoplasm (Fig. 4A, compare middle and right panels). Signal intensity analysis from RNA FISH experiments performed in parallel revealed no differences in CBP80 expression between each condition (Supplementary Fig. 4A and 4B).

**Figure 4:**
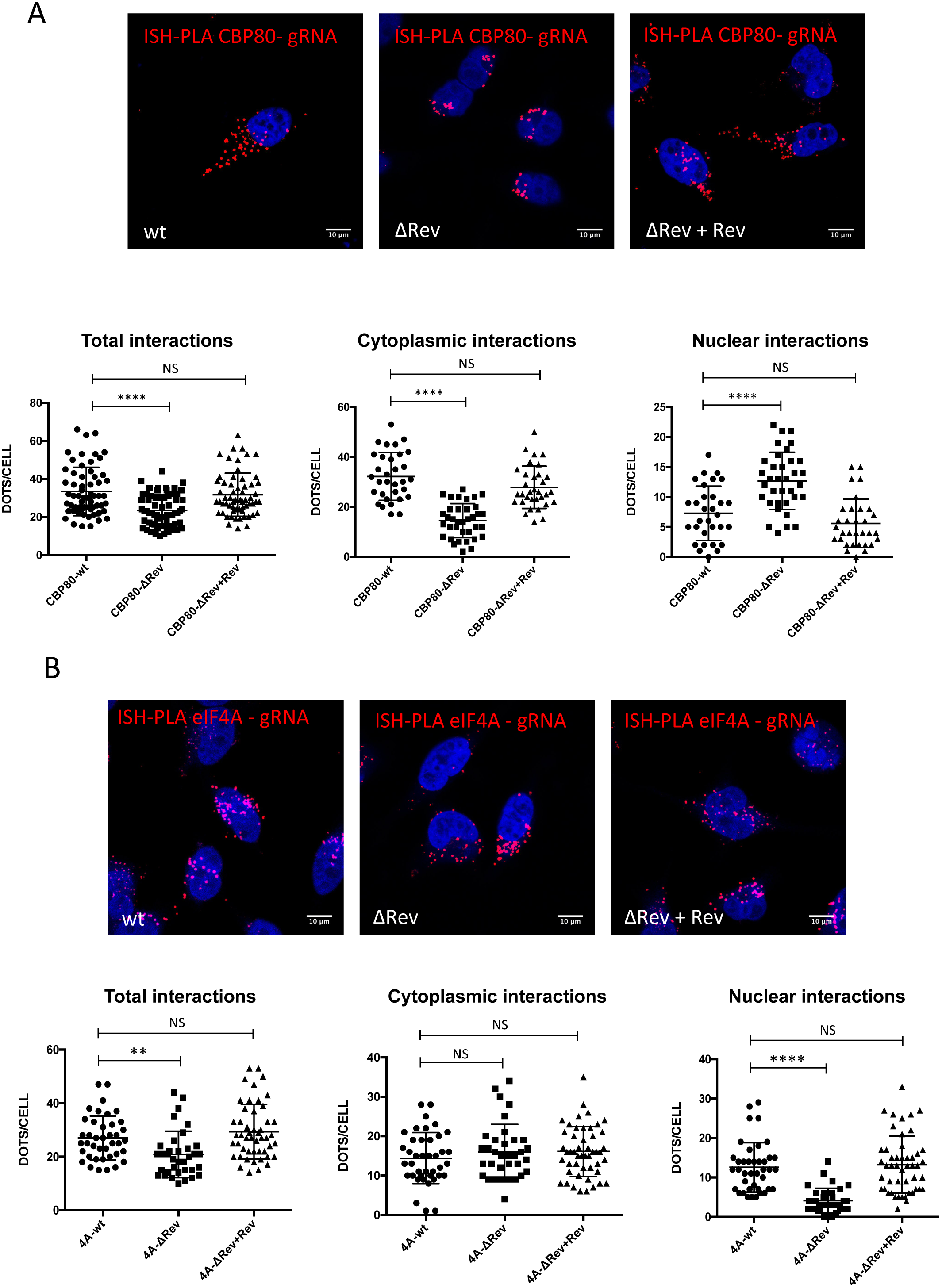
Rev promotes the recruitment of CBP80 and eIF4A to the HIV-1 unspliced mRNA. A) HeLa cells were transfected with 1 μg pNL4.3-wt, 1 μg pNL4.3 ∆Rev or 1 μg pNL4.3 ∆Rev + 0,3 μg pCDNA-Flag-Rev together with 1 μg pCMV-Myc-CBP80. At 24 hpt, the interaction between unspliced mRNA and CBP80 was analyzed by ISH-PLA. Scale bar 10 μm (upper panel). Dots per cell quantifications for total unspliced mRNA-CBP80 interactions (left panel), cytoplasmic interactions (middle panel) and nuclear interactions (right panel) are presented below. All interactions were quantified using ImageJ (^****^*p*<0.0001 and *NS*; non-significant, Mann-Whitney test). B) HeLa cells were transfected with 1 μg of pNL4.3-wt, pNL4.3-∆Rev or 1 μg pNL4.3- ∆Rev + 0,3 μg pCDNA-Flag-Rev together with 1 μg pCIneo-HA-eIF4A. At 24 hpt, the interaction between the unspliced mRNA and eIF4AI was analyzed by ISH-PLA. Scale bar 10 μm (upper panel). Dots per cell for total unspliced mRNA-eIF4A interactions (left panel), cytoplasmic interactions (middle panel) and nuclear interactions (right panel) are presented below. All interactions were quantified using ImageJ (^**^*p*<0.01; ^****^*p*<0.0001 and *NS*; non-significant, Mann-Whitney test) (lower panel)

We also observed that the eIF4AI-unspliced mRNA interactions were reduced in the absence of Rev and restored when the viral protein was expressed *in trans* (Fig. 4B, left panel). Interestingly, we noticed that while most of the interactions between the unspliced mRNA and eIF4AI in the cytoplasm are independent of Rev, the viral protein favors the interaction in the nucleus (Fig. 4B, middle and right panels). Signal intensity analysis from RNA FISH experiments performed in parallel revealed no differences in eIF4A expression between each condition (Supplementary Fig. 4C and 4D). This observation is consistent with our data presented in Fig. 3C in which ectopic expression of eIF4AI favors the accumulation of the unspliced mRNA in the cytoplasm in the presence of Rev and promotes translation in the ΔRev provirus. Interestingly, we observed an increase in the nuclear signal of HA-eIF4AI with the wild type provirus that was absent with the ΔRev provirus suggesting that a fraction of the RNA helicase might translocate to the nucleus in the presence of Rev (Supplementary Fig. 4C and 4E).

Together, these results suggest that Rev regulates the association of CBP80 and eIF4AI with the unspliced mRNA in the cytoplasm and the nucleus, respectively.

### Assembly of a Rev-CBP80-eIF4A complex onto the Rev response element

From our data presented above, it appears that Rev regulates Gag synthesis by associating with CBP80 and eIF4AI and regulating the association of these cellular proteins with the unspliced mRNA in the nucleus and the cytoplasm. Although it was proposed that eIF4AI could be part of the CBC-associated mRNP (72), to our knowledge, the interaction between CBP80 and eIF4AI has never been formally demonstrated. Thus, we finally sought to determine whether CBP80 and eIF4AI interact in cells and whether Rev was influencing such an interaction. As a first approach, we performed PLA to identify and characterize the CBP80-eIF4A interaction and observed that both proteins interact at the nuclear periphery and at a lesser extent in the nucleus (Fig. 5A, left panel). Interestingly, our PLA analysis showed increased interactions in the nucleus when Rev is present (Fig. 5A, right panel). Dots per cell quantification revealed that Rev indeed stimulates the association between CBP80 and eIF4AI (Fig. 5B).

**Figure 5:**
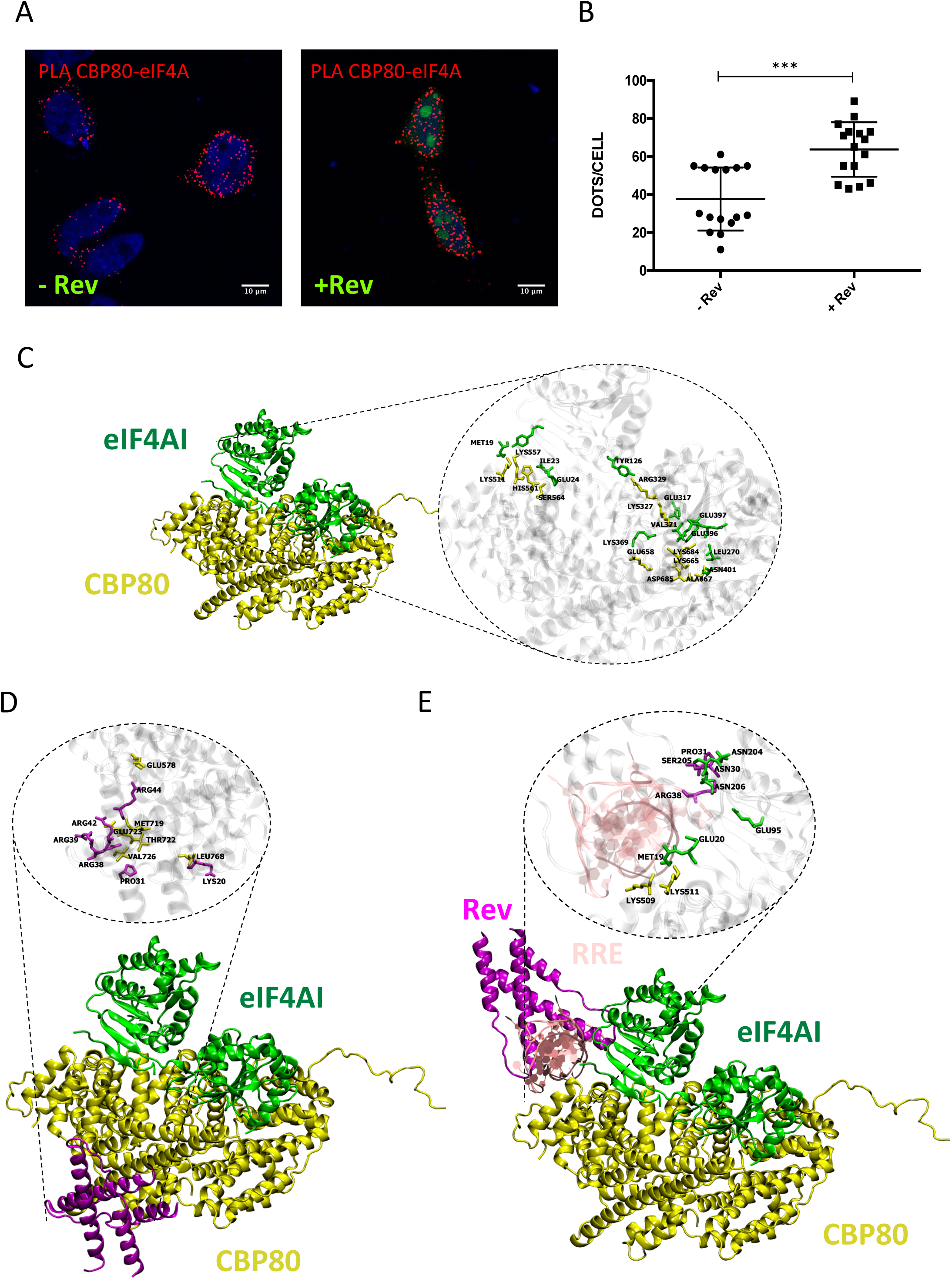
A Rev-CBP80-eIF4AI assembles around the RRE. A) HeLa cells were transfected with 1 μg pCMV-myc-CBP80 and 1 μg pCIneo-HA-eIF4A together 1 μg pEGFP-Rev (pEGFP was used as a control). At 24 hpt, the interaction between HA-eIF4AI and myc-CBP80 was analyzed by PLA. Scale bar 10 μm. B) Dots per cell quantification for eIF4A and CBP80 interactions using ImageJ (^****^*p*<0.0001, Mann-Whitney test). C) Model of the CBP80 (yellow) and eIF4AI (green) complex showing the interaction interface and the residues involved. D) Model of the Rev dimer (magenta), CBP80 (yellow) and eIF4AI (green) complex. The interaction interface and the residues involved in the Rev-CBP80 interaction are presented. E) Model of the Rev dimer (magenta)/RRE (pink), CBP80 (yellow) and eIF4AI (green) complex showing the interaction interface and the residues involved (residues involved in the Rev-RRE interaction were omitted for simplicity).

Having determined that Rev interacts with CBP80 and eIF4AI and that both cellular proteins also interact in cells, we then wanted to gain insights into the potential assembly of a trimeric Rev-CBP80-eIF4AI complex. We performed molecular docking in order to study the non-covalent interactions formed between CBP80 and eIF4AI using previously described structures (51,52). We selected the complex presenting the best binding energy and observed fifteen non-covalent interactions between both cellular proteins (Fig. 5C and Supplementary Table 1). Interestingly, we observed that the eIF4AI binding site on CBP80 overlaps with the CBP20-binding site suggesting that eIF4AI is recruited into a complex alternative to the canonical CBC (Supplementary Fig. 5A and see discussion).

We then used a previously described structure of a Rev dimer in the RNA-free state (73), to investigate whether the viral protein was recruited to the CBP80-eIF4AI complex. Consistent with the lack of non-covalent interactions detected between eIF4AI and the Rev dimer (data not shown), we observed that Rev binds preferentially to CBP80 in the context of our modeled CBP80-eIF4AI complex (Fig. 5D). We mostly observed electrostatic and hydrophobic interactions between positively charged residues in one of the monomers of Rev and negative residues in CBP80 (Supplementary Table 2). Interestingly, residues R38, R39 and R44 in one of the Rev monomers and R44 in the second Rev monomer, which we detected as involved in CBP80 binding (Fig. 5D and Supplementary Table 2), are also involved in RNA binding suggesting that the Rev dimer would not be able to bind the RRE in the context of the Rev-CBP80-eIF4AI complex. However, it was also reported that the interface of the Rev dimer is reorganized upon RNA binding with crossing angles of 120ΰ in the RNA-free state and 50° in the RNA-bound state (55,73). Thus, we finally sought to evaluate whether RNA binding by Rev has an impact in the assembly of the Rev-CBP80-eIF4AI complex. For this, we performed docking analyses using the structure of the Rev/RRE complex and our modeled CBP80-eIF4AI complex. Interestingly, we observed that the presence of the RRE results in the complete reorganization of the trimeric complex (Fig. 5E and Supplementary Table 3). As such, we observed that both CBP80 and eIF4AI establish non-covalent interactions with the RRE. In addition, while the interactions between Rev and CBP80 were lost, we observed that eIF4AI was able to perform contacts with the viral protein in its RNA-bound state (Fig. 5E and Supplementary Table 3). Taking together, these data suggest that Rev forms a complex with CBP80 and eIF4AI that reorganizes when the viral protein binds to the RRE. This complex drives Gag synthesis from the unspliced mRNA.

## Discussion

Gag synthesis from the unspliced mRNA is a critical step during HIV-1 replication necessary for the efficient production of the viral progeny. Indeed, stoichiometric studies revealed that up to 5000 molecules of Gag are required to build one single viral particle (74), indicating that the unspliced mRNA needs to be efficiently expressed. However, since this viral mRNA contains functional introns, it must overcome surveillance mechanisms that induce its nuclear retention and degradation before it reaches the translational machinery in the cytoplasm to produce Gag (75). As such, understanding how the unspliced mRNA is efficiently exported from the nucleus and translated in the cytoplasm is not only critical to improve our knowledge on the molecular mechanisms driving viral gene expression but is also important to identify new pathways and/or interactions occurring within the cell or induced by the virus that could be targeted by novel antiretroviral drugs.

Although the unspliced mRNA associated to Rev, CRM1 and other co-factors does not resemble to a canonical mRNP that must be directed to the ribosomes upon nuclear export (i.e., an mRNA associated to cellular components such as TREX, NXF1 and the EJC), translation of the unspliced mRNA is highly efficient in cells and nuclear export across the Rev/CRM1 pathway is critical for ribosome recruitment (37,76). In this study, we provide evidence that the viral protein Rev acts as a nuclear imprint critical for nuclear export and translation of the unspliced mRNA (Fig. 1). We reasoned that recruitment of Rev might serve as a platform for the spatiotemporal recruitment of host factors required to interconnect nuclear export and translation of the unspliced mRNA. In order to study the interaction between the unspliced mRNA and host proteins, we developed the ISH-PLA strategy, which allowed us to identify and quantify unspliced mRNA-protein interactions but also to determine the cellular location in which such interactions occur. In agreement with previous data (69), we observed that the HIV-1 unspliced mRNA is preferentially associated to the CBC subunit CBP80 both in the nucleus and the cytoplasm (Fig. 2). Interestingly, we also confirmed that CBP80 interacts with Rev and showed that the CBC subunit supports the activity of Rev in nuclear export and translation. Despite the fact that our *in silico* data suggest that Rev does not interfere with the interaction between CBP80 and CBP20 (data not shown), it is still unknown whether CBP80 alone, or in the context of the CBC, is responsible of these functions. Indeed, we observed that eIF4AI, which also interacts with CBP80 and Rev, would interfere with recruitment of CBP20. Moreover, our unpublished data also shows that CTIF, the CBP80/20-dependent translation initiation factor is a potent inhibitor of Gag synthesis (Garcia-de-Gracia *et al*. manuscript in preparation). These observations strongly suggest that CBP80 might promote Gag synthesis in the context of a non-canonical CBC. Interestingly, a recent study reported the existence of an alternative CBC formed by CBP80 and NCBP3, a novel cap-binding protein specifically associated to mRNA nuclear export (19). Thus, it would be of interest to determine whether the translating unspliced mRNA is indeed associated to the CBC and which of the cap-binding proteins, CBP20 or NCBP3, is bound to the cap structure of the viral transcript during translation. In this sense, the HIV-1 unspliced mRNA was shown to contain a m^2,2,7^ GpppG trimethylated cap in a process catalyzed by the methyltransferase PIMT and dependent on the presence of Rev (77). Interestingly, increased cap trimethylation by PIMT overexpression was shown to promote Gag synthesis suggesting that a trimethylated cap favors polysome association of the unspliced mRNA (77). Since trimethylation reduces the affinity of CBP20 and eIF4E for the cap (78,79), it would be of interest to test the affinity of NCBP3 for m^2,2,7^ GTP and whether this new cap-binding protein drives translation of the HIV-1 unspliced mRNA together with CBP80. It should be mentioned that our ISH-PLA analyses do not discard an association of the unspliced mRNA with the cap-binding protein eIF4E, which probably reflects the proportion of viral transcripts that contain monomethylated caps. Previous reports have shown that HIV-1 Gag synthesis and replication are maintained under inhibition of eIF4E-driven cap-dependent translation (69,80,81). Whether our Rev-CBP80-eIF4AI complex or the IRES-driven mechanism of ribosome recruitment are responsible of maintaining Gag synthesis under unfavorable conditions needs to be further investigated.

We also identified the DEAD-box RNA helicase eIF4AI as an additional partner of Rev (Fig. 3). Interestingly, we observed that ectopic expression of eIF4AI promoted both the cytoplasmic accumulation of the unspliced mRNA or translation depending on whether Rev was present or not. Consistent with these observations, we observed that Rev promotes the interaction between eIF4AI and the unspliced mRNA in the nucleus (Fig. 4). Since inhibition of eIF4AI/II function by hippuristanol treatment resulted in a strong inhibition of unspliced mRNA translation from the wild type provirus (data not shown), we propose that the RNA helicase plays a dual role during viral replication by assisting Rev during nuclear export and by promoting unspliced mRNA translation independently of the viral protein. Further work is necessary to decipher the mechanism by which eIF4AI cooperates with Rev during nuclear export. We further showed that CBP80 associates with eIF4AI and this interaction was stimulated in the presence of Rev (Fig. 5). Although this interaction was proposed some time ago (72), to our knowledge this is the first experimental evidence for the association between CBP80 and eIF4AI. Since we observed that the CBP20 and eIF4AI might compete for binding CBP80, it would be of interest to determine the consequences of the CBP80-eIF4AI interaction, for example, during the pioneer round of translation or NMD. It would also be interesting to evaluate whether Rev is able to regulate these cellular processes. Interestingly, our molecular docking analyses using the Rev dimer bound to the RRE suggest that the viral RNA serves as a platform for the assembly of the trimeric Rev-CBP80-eIF4AI complex. Thus, we propose a model in which Rev interconnects CRM1-dependent nuclear export with ribosome recruitment of the unspliced mRNA by driving the recruitment of host factors required for both processes (Fig. 6). The proper and timely assembly of such a Rev-dependent mRNP will determine the efficient association of the unspliced mRNA with the host machineries for nuclear export and translation initiation. Last but not least, the small molecule ABX464, currently under a phase II clinical trial, was shown to interfere with the Rev-CBP80 interaction (82,83). Therefore, results presented here will be useful either for the better understanding of the mechanism of action of this small molecule or for the rational design of new drugs targeting the Rev-CBP80-eIF4A complex.

**Figure 6:**
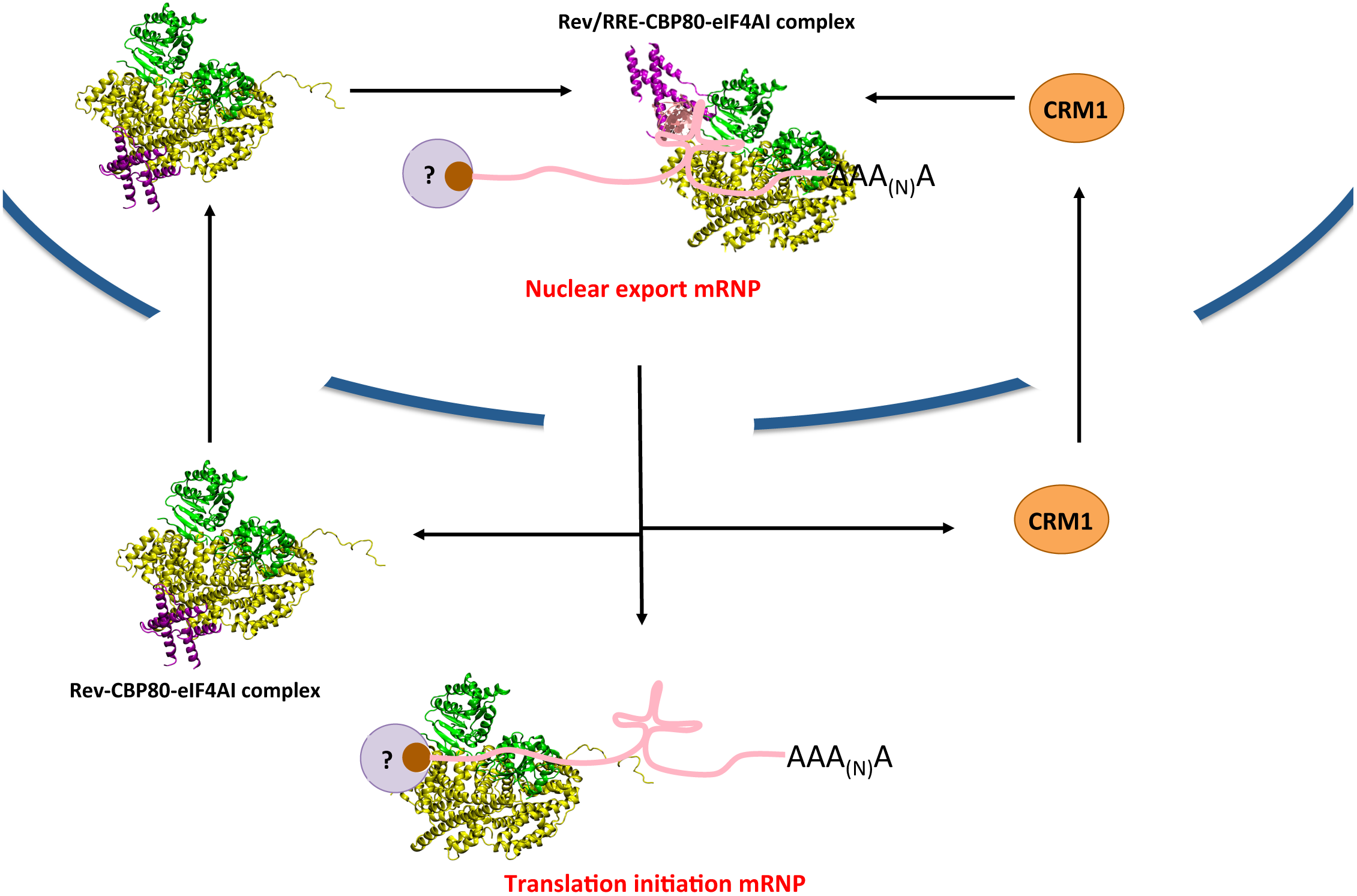
A model for the function of the Rev-CBP80-eIFAI complex on Gag synthesis. The Rev-CBP80-eIF4AI complex might assemble into the cytoplasm and be imported to the nucleus to be recruited onto the RRE. Once Rev binds to CRM1 through its NES, this nuclear export mRNP translocates to the cytoplasm. Upon dissociation of Rev and CRM1, CBP80 remain associated to the gRNA to promote translation in a Rev-dependent manner. In contrast, eIF4AI is either recruited or remain associated to the gRNA independently of Rev in order to promote gRNA translation. The cap-binding protein associated to the HIV-1 gRNA is still unknown.

## Acknowledgements

Authors wish thank to Dr. Richard Cerione (Cornell University, USA), Dr. Jonathan Karn (Case Western Reserve University, USA), Dr. Yoon Ki Kim (University of Korea) and Dr. Marcelo López-Lastra (PUC, Chile) for their generous provisions of plasmids and reagents. Authors wish also thank to Alessandra Dellarossa for technical support. The following reagents were obtained through the NIH AIDS Reagents Program, Division of AIDS, NIAID, NIH: HIV-1 p24 Monoclonal Antibody (183-H12-5C) from Dr. Bruce Chesebro and Kathy Wehrly, THP-1_ATCC_ from Drs Li Wu and Vineet N. Kewal Ramani and H9 cells from Dr. Robert Gallo.

